# Rewired signaling network in T cells expressing the chimeric antigen receptor (CAR)

**DOI:** 10.1101/2019.12.19.883496

**Authors:** Rui Dong, Kendra A. Libby, Franziska Blaeschke, Alexander Marson, Ronald D. Vale, Xiaolei Su

## Abstract

The chimeric antigen receptor (CAR) directs T cells to target and kill specific cancer cells. Despite the success of CAR T therapy in clinics, the intracellular signaling pathways that lead to CAR T cell activation remain unclear. Using CD19 CAR as a model, we report that, similar to the endogenous T cell receptor (TCR), antigen-engagement triggers the formation of CAR microclusters that transduce downstream signaling. However, CAR microclusters do not coalesce into a stable central supramolecular activation cluster (cSMAC). Moreover, LAT, an essential scaffold protein for TCR signaling, is not required for microcluster formation, immunological synapse formation, and actin remodeling following CAR activation. Meanwhile, CAR T cells still require LAT for the normal production of the cytokine IL-2. Together, these data show that CAR T cells can bypass LAT for a subset of downstream signaling outputs, thus revealing a rewired signaling pathway as compared to native T cells.

## Introduction

The development of T cells engineered with chimeric antigen receptors (CARs) has launched a new era of cell-based therapy for cancer. CAR is a synthetic single-pass transmembrane receptor that is intended to mimic the signal transducing function of the T cell receptor (TCR). CAR recognizes a tumor antigen and activates pathways leading to T cell activation. CAR-T cell therapy, as approved by FDA, has been successfully applied to treat patients with B cell cancers including acute lymphoblastic leukemia and non-Hodgkin lymphoma^1–3^. However, many patients do not respond to CAR T therapy and some suffer from severe neurotoxicity and cytokine release syndrome^4,5^. Furthermore, there are unexplained aspects of the CAR T cell response to antigen stimulation. CARs, for example, trigger faster tyrosine phosphorylation of downstream effectors and faster cytotoxic granule release in CD8^+^ T cells^6^. Perhaps even more surprisingly, CAR-CD4^+^ T cells can kill target cells as efficiently as CAR-CD8^+^ T cells, although the basis of this phenomenon is not known^7^. A better understanding of the molecular mechanism of CAR signaling is needed for explaining the unique aspects of CAR T cells and could suggest new strategies for designing better CAR-T therapies targeting broad types of cancers.

Here, we sought to define the signaling pathways downstream CAR activation in both immortalized human T cell lines (Jurkat T cells) and primary human T cells. We found that, similar to TCR, CARs form near micron-sized clusters that serve as signaling platforms to transduce the activation signal from antigen engagement. However, LAT, an adaptor protein essential for TCR microcluster formation, is dispensable for the formation of CAR microclusters and CAR-induced immunological synapses. CAR also can bypass LAT to activate LAT’s downstream signaling pathways leading to actin reorganization and synapse formation. Together, these data suggest that CAR T cells display a rewired signaling pathway as compared to the TCR pathway in native T cells.

## Results and Discussion

### CARs form microclusters upon ligand engagement

We used the 3^rd^ generation CD19-CAR as a model, which is activated by the CD19 molecule on the B cell surface. A transmembrane domain derived from CD8α connects the extracellular single chain variable fragment (scFv) recognizing CD19 to the intracellular domain composed of the cytoplasmic part of TCR subunit CD3ζ, co-receptor CD28, and 4-1BB (**Figure 1A**). A GFP tag was introduced at the C-terminus to visualize CAR dynamics. GFP-tagged CAR was stably introduced into Jurkat T cells by lentiviral transduction. To stimulate these CAR-T cells, we used a supported lipid bilayer-based activation system^8^, which is compatible with high signal-to-noise imaging of the T cell plasma membrane by total internal reflection fluorescence (TIRF) microscopy. In this system, recombinant CD19 proteins, the ligand for CAR, were attached to the supported lipid bilayers, as a surrogate for antigen-presenting cells. To facilitate cell adhesion, the supported lipid bilayer also was coated with ICAM-1, an integrin ligand **(Figure 1B**).

**Figure 1.**
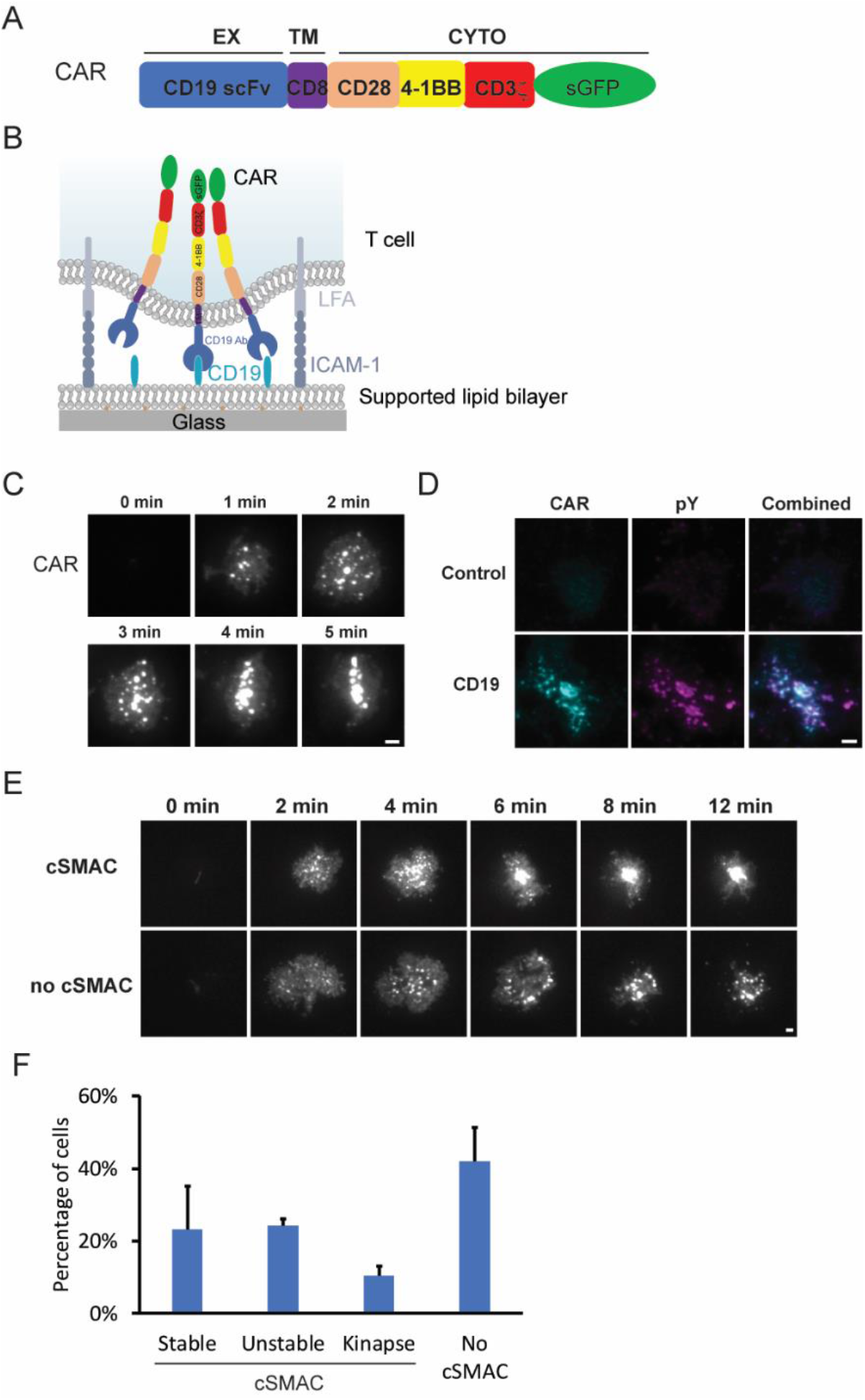
CAR forms microclusters that are signaling-active. **A.** Domain structure of CAR. **B.** Schematic of the assay for imaging CAR dynamics. **C.** Time courses of microcluster formation of CAR upon CD19 engagement. **D.** Enrichment of phosphortyrosine (pY) at CAR microclusters. **E.** CAR microclusters underwent centripetal movement to form cSMAC (upper panel) or remained separated (lower panel). Scale bar: 2 μm. **F.** Quantification of the categories regarding the formation of CAR cSMAC. Shown are the mean + SD. *n*=3 independent experiments. 38 to 53 cells were quantified in each experiment.

When CAR-T cells were plated onto the CD19- and ICAM-1-coated bilayers, CAR microclusters rapidly formed as cells spread on the supported lipid bilayer (**Figure 1C**). These CAR microclusters were enriched in phosphotyrosines, indicating that they are spots of active signaling (**Figure 1D**). In contrast, in the absence of CD19, neither microclusters nor phosphotyrosine signals were present on the T cell membranes (**Figure 1D**). These data show the CD19 binding to CAR triggers the formation of signaling-competent microclusters, which is very similar to what has been reported for pMHC-triggered TCR microclusters^9^,^10^.

We then monitored the behavior of CAR microclusters in a longer term. In about one-quarter of the CAR-T cells, CAR microclusters translocated to the cell center and formed a stable disk-like structure (**Figure 1E**, upper panel and **Movie S1**), which resembles the central supramolecular activation cluster (cSMAC)^11^. However, in another quarter of cells, the CAR cSMAC initially formed but then quickly disassembled (**Movie S2**). The average time between cell landing and cSMAC disassembly was ~7 min. We also observed moving synapses (kinapse) in about 11% of cells (**Movie S3**). Most strikingly, in ~40% of cells, individual microclusters remained separated and did not form a cSMAC (**Figure 1E** lower panel, **Movie S4**). Together, these data (**Figure 1F**) suggest that CARs form microclusters, but these generally do not coalesce into a stable cSMAC.

### CARs can bypass LAT to form microclusters and induce immunological synapse formation

Next, we determined the molecular requirements for transducing CAR signaling. LAT, or linker for activation of T cells, is an essential adaptor for TCR signaling^12^. Deletion or mutation of LAT impairs formation of T cell microclusters^13^,^14^ as well as signal transduction to downstream pathways including calcium influx, MAPK activation, IL-2 production, and actin remodeling^15,16^. Firstly, we reconfirmed that TCR microclusters did not form in a LAT-deficient Jcam2.5 T cell line (**Figure 2A**, left panel). Surprisingly, however, we found that CARs still formed microclusters in the same LAT-deficient line (**Figure 2A** and **2B**).

**Figure 2.**
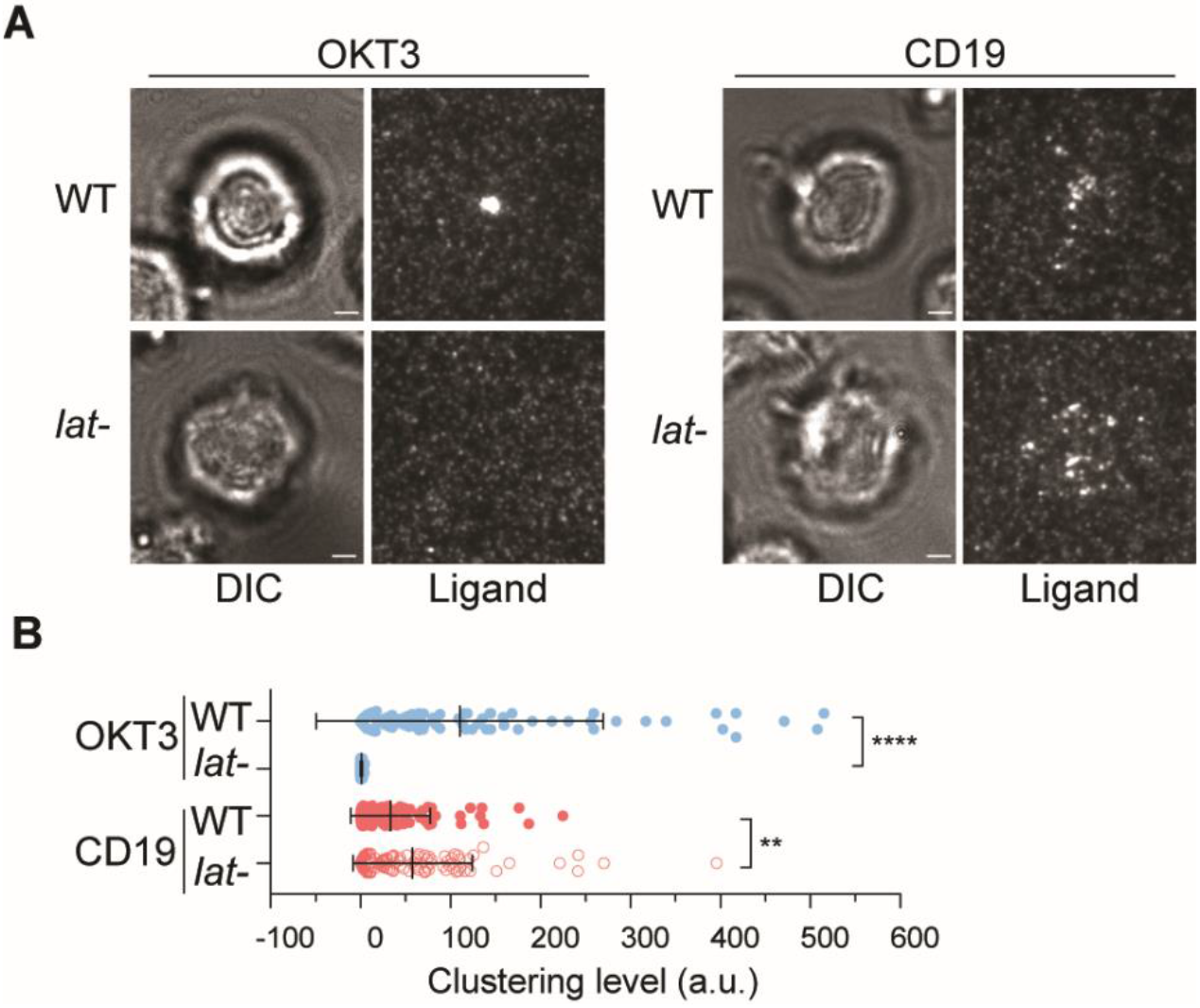
LAT is not required for CAR-induced microcluster formation. **A.** WT or LAT-deficient Jurkat cells expressing CAR stimulated on planar lipid bilayers coated with OKT3 (anti-TCR antibody) or CD19 (CAR antigen). TIRF microscopy revealed clustering of Ax647–labeled streptavidin-Biotin-OKT3 or CD19, which serves as a probe for TCR or CAR respectively. Scale bar, 2 μm. **B.** Quantification of clustering as normalized variance^13^. *n* = 100 cells. Shown are the mean ± SD. **** *p* < 0.0001. ** *p* = 0.0022.

Next, we investigated whether CAR microclusters can recruit signaling components downstream of LAT. Gads is a cytosolic adaptor that is, after TCR activation, recruited to the plasma membrane by LAT. Gads bridges LAT to SLP76, which further connects the LAT microclusters to the actin polymerization machinery^9^. We found that Gads could be recruited to CAR microclusters in both the presence and absence of LAT. The amount of Gads relative to CAR recruited to membranes in LAT-deficient cells is comparable to that in the wild-type cells (**Figure 3A**). This result indicates Gads can still be recruited to CAR microclusters without LAT.

**Figure 3.**
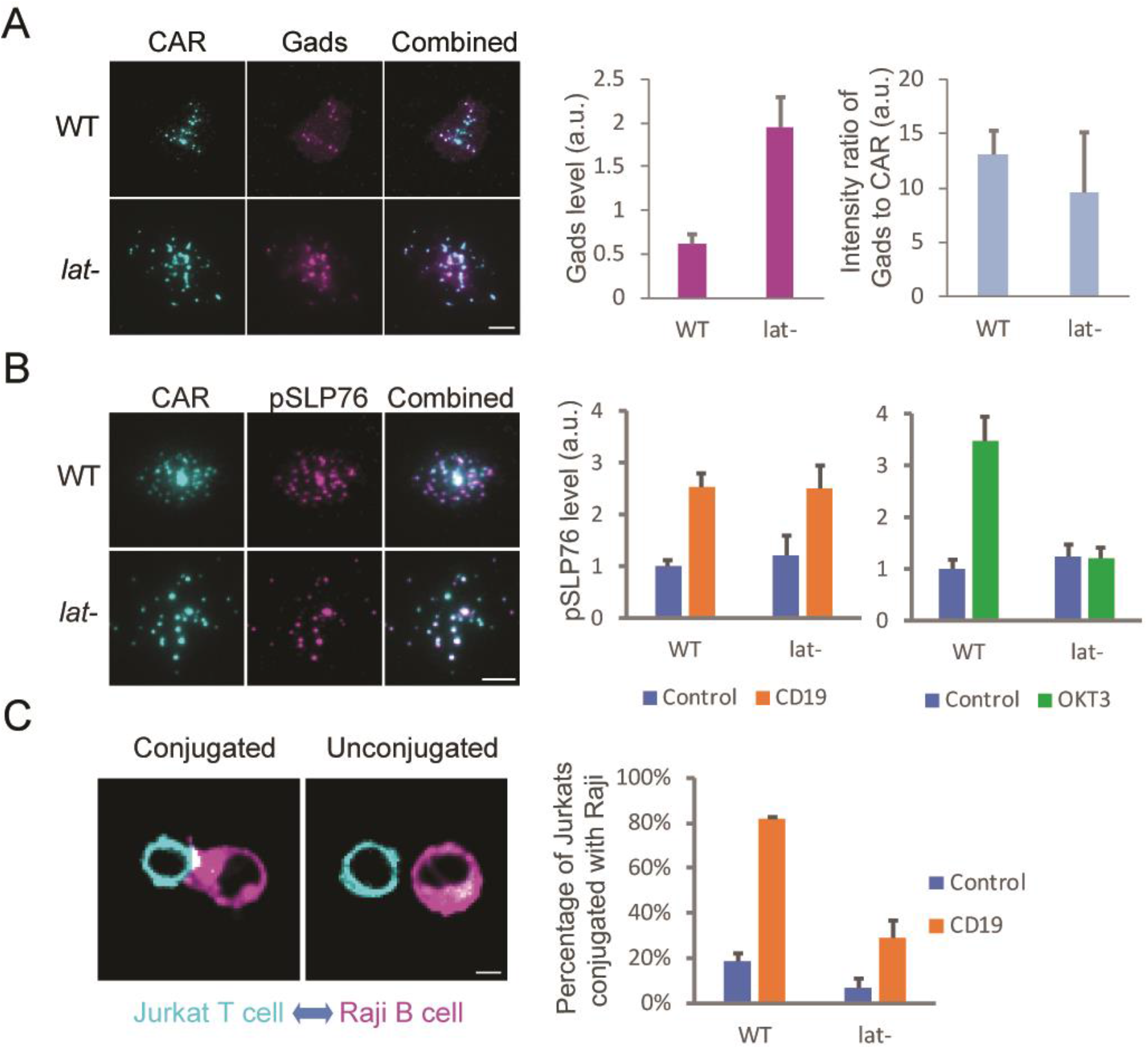
CAR microclusters can activate Gads-SLP76 pathway in the absence of LAT. **A.** Gads is recruited to CAR microclusters in LAT-deficient cells. N=33 or 34 cells. Shown are the means ± SE. **B.** Phosphorylated SLP76 is enriched in CAR microclusters in both wild-type and LAT cells. *n* = 50 cells. Shown are the means ± SE. **C.** LAT is dispensable for cell conjugates formation between Jurkat T cells expressing CAR and Raji B cells expressing CD19. Jurkat cells are mixed with Raji cells at a 1:1 ratio. K562 cells, which do not express CD19, serve as a negative control. Three independent experiments were performed and over 100 cells were scored in each experiment. Shown are the means ± SD. Scale bar: 5 μm.

We then investigated whether LAT is required for the activation of SLP76. SLP76 is a constitutive binding partner for Gads^17^ and activated SLP76 recruits Nck, an important regulator of actin assembly^18^, to LAT clusters on the membrane. To assess the activation level of membrane-associated SLP76, we performed immunofluorescence microscopy using a phospho-SLP76 antibody. We found that pSLP76 levels at the membrane were similar between the wild-type and LAT-deficient cells following CD19-dependent CAR activation (**Figure 3B**). In contrast, the level of pSLP76 was not increased by OKT3, the activating antibody for TCR, in LAT-deficient cells (**Figure 3B**). These results suggest that LAT is required for TCR-triggered, but not CAR-triggered activation of SLP76. In support of the idea that LAT is not required for Gads-SLP76-actin pathway, in LAT-deficient CAR-T cells, CAR microclusters were still robustly translocated from the periphery to the cell center (**Movie S5**). This result indicates that CAR engagement with CD19 induces actin polymerization-dependent retrograde flow in LAT knockout cells.

Actin polymerization also stabilizes immunological synapse formation between a T cell and an antigen-presenting cell. To test if LAT is required for CAR-T cell synapse formation, we incubated CAR-T cells with Raji B cells that express CD19. We found LAT-deficient cells can still form synapses with Raji B cells, though the efficiency is lower than the wild-type cells (**Figure 3C**). This result suggests that LAT promotes but is not essential for CAR-induced immunological synapse formation. Together, these data suggest that LAT is not essential for transducing signaling from CAR to the actin polymerization pathway.

### LAT is required for CAR-triggered PIP2 hydrolysis and IL-2 production

In parallel, we determined if LAT is required for PIP2 signaling following CAR activation. We used mCherry-labeled C1 domain as a reporter for diacylglycerol (DAG)^19^, the product of PIP_2_ hydrolysis. In contrast to wild-type cells, in which the C1 domain was robustly recruited to the plasma membrane upon CAR activation, C1 was not recruited in LAT-deficient cells (**Figure S1A**). Consistent with a defect in PIP_2_ hydrolysis, CAR also did not induce IL-2 secretion in LAT-deficient cells (**Figure S1B**). Together, these data suggest LAT is essential for CAR-triggered PIP2 signaling and IL-2 production in Jurkat T cells.

### LAT is not required for CAR signaling in human primary T cells

Next, we sought to determine if a LAT-independent CAR signaling pathway exists in primary T cells. To do so, we knocked out the LAT gene in human primary T cells by CRISPR-based genome targeting. The LAT protein levels were reduced in the overall cell population to 12% as revealed by immunoblot analysis (**Figure 4A**). CAR-GFP was then introduced into wild-type or LAT null T cells by lentiviral transduction. TIRF microscopy revealed that TCR microclusters were formed in wild-type but not LAT null cells following OKT3 engagement on planar lipid bilayers. In contrast, CAR microclusters were formed in both wild-type and LAT null cells following CD19 engagement (**Figure 4B**). Moreover, SLP76, the adaptor linking CAR to actin, was equally activated in wild-type and LAT null CAR T cells, as determined by a phospho-SLP76 antibody (**Figure 4C**). Interestingly, we also found that LAT null CAR T cells produced IL-2, though at a lower level than wild-type CAR T cells (**Figure 4D**). This result differs from that obtained with Jurkat T cells, in which IL-2 production is completely abolished in LAT-deficient cells (**Figure S1B**). The mechanism underlying this difference is unclear, potentially due to the different signaling responses between Jurkat and primary T cells^20^. Together, these data suggest that LAT is not required for CAR microcluster formation and SLP76 activation in human primary T cells.

**Figure 4.**
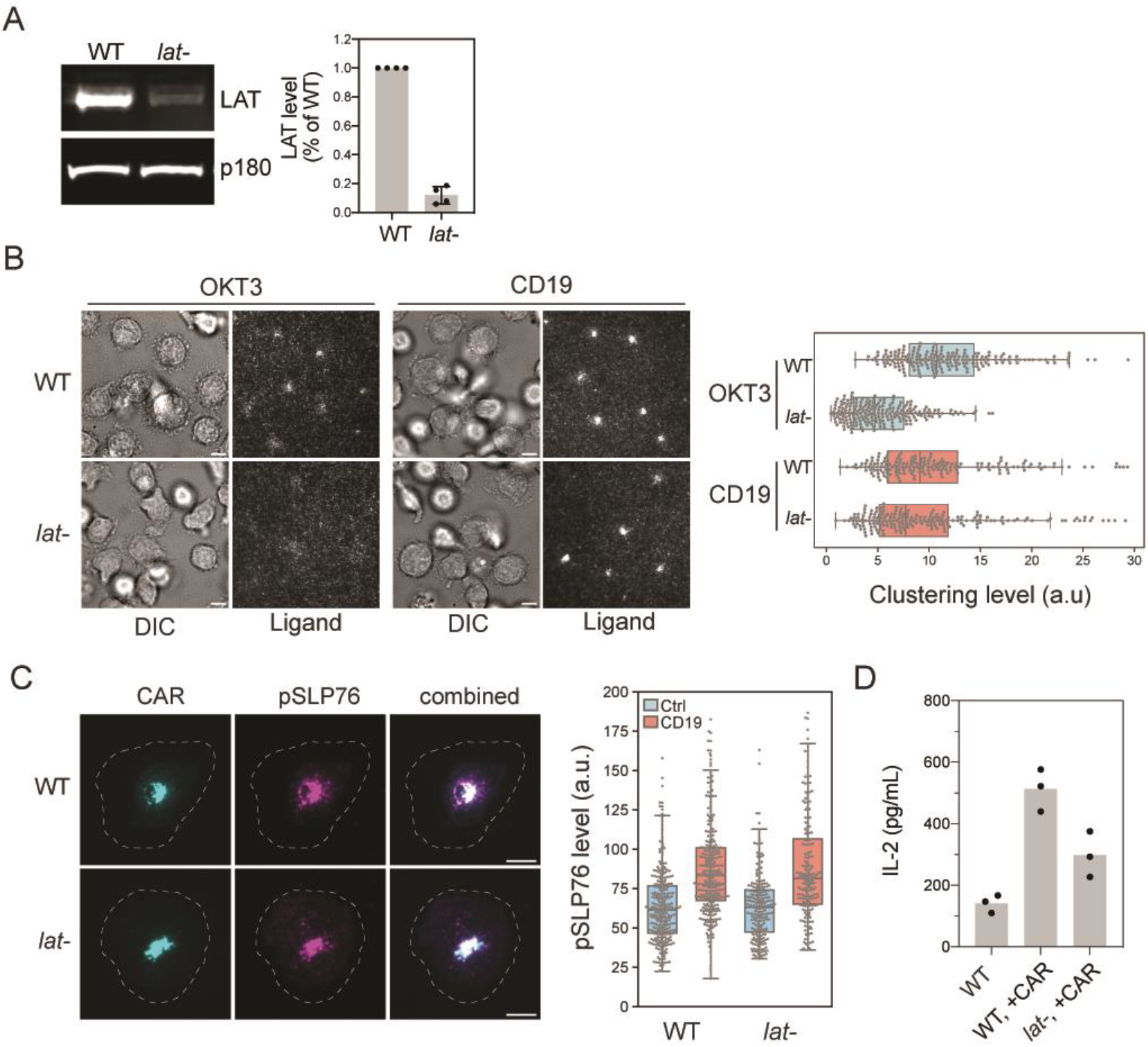
LAT is not required for CAR microcluster formation and SLP76 activation in human primary T cells. **A.** Western blot showed that LAT is efficiently depleted in human primary T cells using CRISPR. Shown are the means ± SD. **B.** WT or LAT-deficient CAR T cells were stimulated on planar lipid bilayers coated with biotin-OKT3 (left) or biotin-CD19 (right). TIRF microscopy revealed clustering of Alexa647–labeled streptavidin-Biotin-CD19. **C.** Phosphorylated SLP76 is enriched in CAR microclusters in both WT and lat-T cells. As a control, pSLP76 levels are significantly lower on bilayers without CD19. For Quantification of pSLP76 in **B** and **C**, *n* > 200 cells were scored, and the cells were isolated from two independent donors. Shown are the means ± SD. Clustering is quantified as normalized variance. Scale bar, 5 μm. **D.** T cells were incubated with Raji B cells at a 4:1 ratio and IL-2 were measured by ELISA at 24hr. Data were pooled from three donors.

### A model for CAR T signaling pathway

Our data reveal that CAR exhibits differences in its signaling mechanism from TCR. LAT plays a crucial role in signaling after TCR engagement. Tyrosine-phosphorylated LAT recruits several molecules, including Grb2, PLCγ1, and Gads, which promote microcluster formation. Microclusters further transduce signal to downstream pathways that promote synapse formation and cytokine production (**Figure 5A**). In contrast, when CAR is engaged, it can induce microcluster formation even in the absence of LAT. CAR could directly recruit LAT binding partner Gads, to trigger downstream signaling that leads to actin remodeling and immunological synapse formation. However, LAT is still partially required for CAR-induced cytokine production (**Figure 5B**).

**Figure 5.**
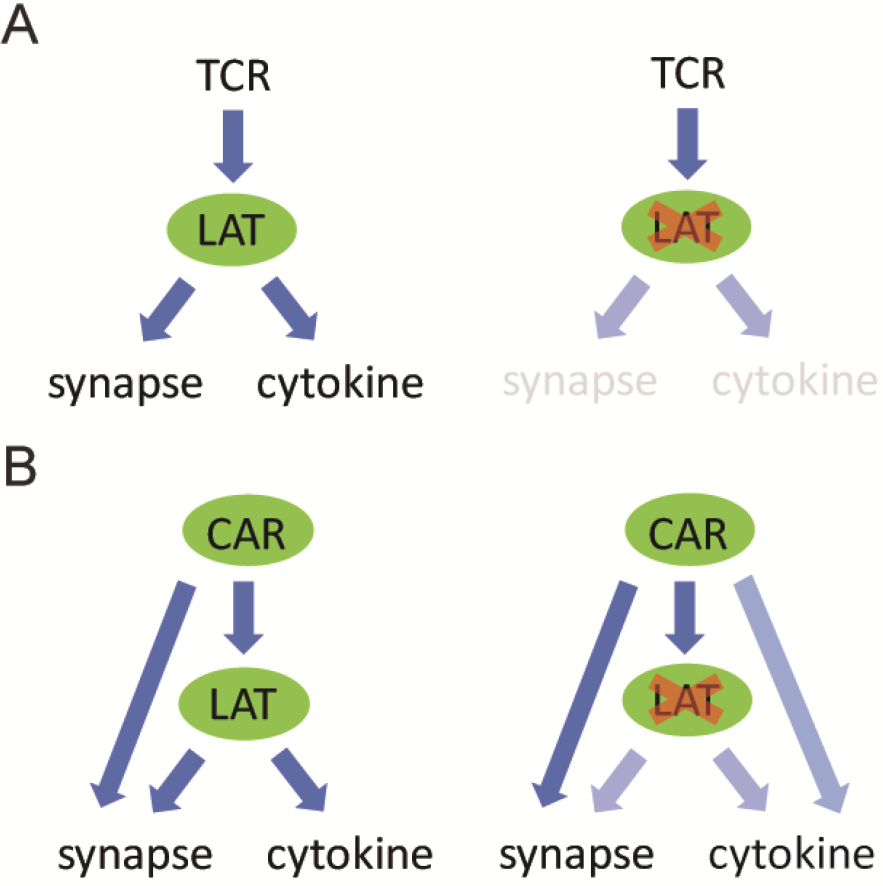
A model comparing TCR-versus CAR-induced signaling. **A.** Activated TCR triggers LAT phosphorylation, which promotes microcluster formation. Microclusters further transduce signal to downstream pathways that induce synapse formation and cytokine production. No microclusters or synapses are formed in the absence of LAT. **B.** In contrast, when CAR is activated, it induces microcluster formation even in the absence of LAT. CAR could directly recruit LAT binding partners to trigger downstream signaling that leads to synapse formation and partial cytokine production.

The discovery of a LAT-independent signaling pathway may be related to and provide an explanation for previous reports showing that CAR triggers faster proximal signaling and faster killing of target cells than TCR^6^. CAR could take a shortcut to bypass LAT, and directly recruit downstream signaling proteins Grb2, Gads, and PLCγ1 to transduce signaling. The multivalent interactions between CAR and SH2 domain-containing adaptor proteins might drive the formation of CAR microclusters, even in the absence of LAT. However, additional work will be needed to dissect more details on the LAT-independent signaling and how that changes the overall kinetics of the CAR T cell signaling response.

## Supporting information

Movie S1. Stable cSMAC

Movie S2. Unstable cSMAC

Movie S3. Kinapse

Movie S4. No cSMAC

Movie S5. Clusters in lat- cells

## Acknowledgement

R.D. was supported by a Jane Coffin Childs Postdoctoral Fellowship. K. L. was supported by a Yale College First-Year Summer Research Fellowship in the Sciences and Engineering. F. B. was supported by the Care-for-Rare Foundation, the German Research Foundation (DFG), and an NIH funding for the HIV Accessory & Regulatory Complexes (HARC) Center (P50 GM082250). A.M. holds a Career Award for Medical Scientists from the Burroughs Wellcome Fund, is an investigator at the Chan Zuckerberg Biohub and has received funding from the Innovative Genomics Institute (IGI) and the Parker Institute for Cancer Immunotherapy (PICI). R.D. is an investigator of the Howard Hughes Medical Institute. X.S. has received support from the American Cancer Society Institutional Research Grant, the Charles H. Hood Foundation Child Health Research Awards, the Andrew McDonough B+ Foundation Research Grant, the Gilead Sciences Research Scholars Program in Hematology/Oncology, and the Rally Foundation a Collaborative Pediatric Cancer Research Awards Program.

## Competing Financial Interests

The authors declare competing financial interests: A.M. is a co-founder of Arsenal Biosciences and Spotlight Therapeutics. A.M. serves as on the scientific advisory board of PACT Pharma, is an advisor to Trizell and was a former advisor to Juno Therapeutics. The Marson Laboratory has received sponsored research support from Juno Therapeutics, Epinomics, Sanofi and a gift from Gilead.

**Figure S1.**
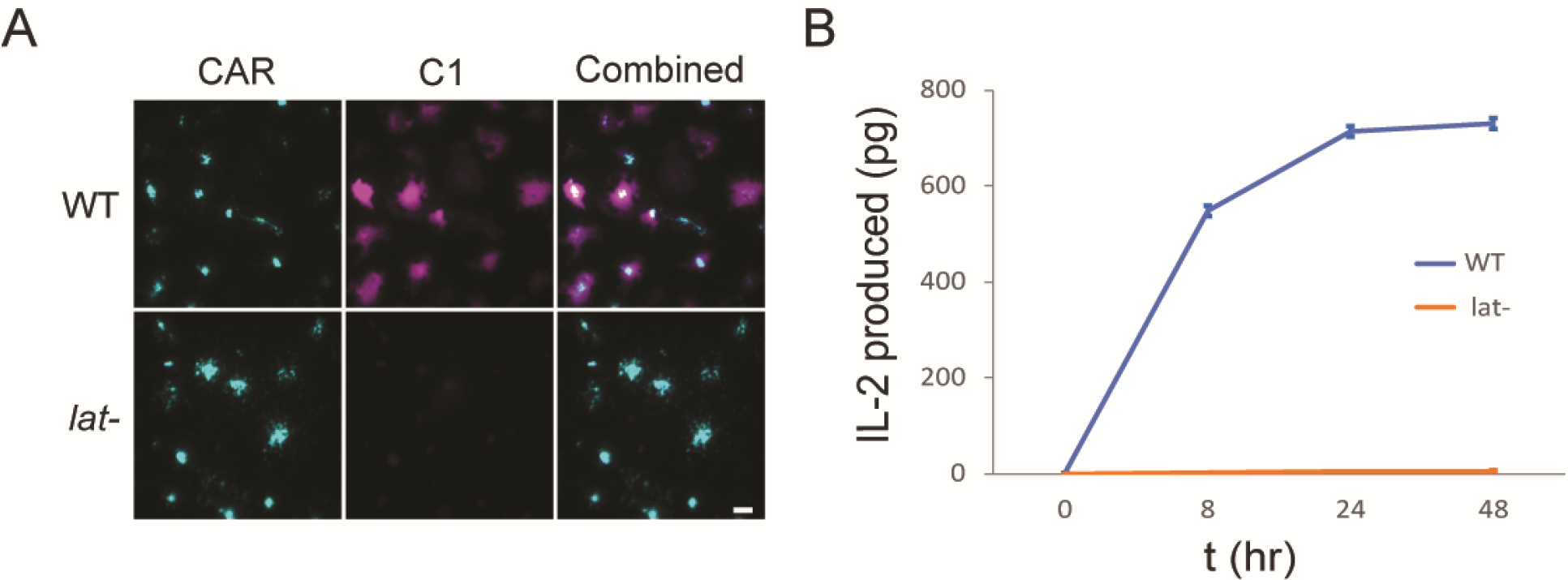
LAT is required for PIP2 hydrolysis and IL-2 production in Jurkat T cells. **A.** LAT is required for PIP2 hydrolysis in CAR-induced signaling. PIP2 is hydrolyzed to generate DAG and IP3, following the receptor activation. The DAG reporter C1 is recruited to plasma membrane only in wild-type but not LAT deficient cells. Scale bar: 5 μm. **B.** LAT is required for IL-2 production. CAR-T cells were mixed with Raji B cells expressing CD19 for 24 hrs. IL-2 produced was measured by ELISA. Shown are means ± SD. *n* = 3 independent experiments.

### Supplementary Movies

**Movie S1. CAR forms a stable cSMAC.** TIRF microscopy revealed an example showing CAR-GFP forming a stable cSMAC as the cell lands on supported lipid bilayers coated with CD19 and ICAM-1. Shown is a field view of 23 μm x 23 μm.

**Movie S2. CAR forms a cSMAC that disassembles early.** TIRF microscopy revealed an example showing CAR-GFP forming an unstable cSMAC that quickly disassembles as the cell lands on supported lipid bilayers coated with CD19 and ICAM-1. Shown is a field view of 23 μm x 23 μm.

**Movie S3. CAR forms a moving cSMAC.** TIRF microscopy revealed an example showing CAR-GFP forming a moving cSMAC (kinapse) as the cell lands on supported lipid bilayers coated with CD19 and ICAM-1. Shown is a field view of 23 μm x 23 μm.

**Movie S4. CAR forms a scattered pattern.** TIRF microscopy revealed an example showing CAR-GFP remaining scattered and not coalescing into a cSMAC as the cell lands on supported lipid bilayers coated with CD19 and ICAM-1. Shown is a field view of 23 μm x 23 μm.

Movie S5. CAR forms a moving cSMAC. TIRF microscopy revealed an example showing CAR-GFP microclusters undergoing centripetal movements as the LAT-deficient cell lands on supported lipid bilayers coated with CD19 and ICAM-1. Shown is a field view of 25 μm x 25 μm.

## Methods

### Key Resources Table

| Reagent type | Designation | Source or reference | Additional information |
| --- | --- | --- | --- |
| Cell line | Jurkat T cell E6.1 | UCSF Cell Culture Facility |  |
| Cell line | Jurkat T cell Jcam2.5 LAT deficient | UCSF Cell Culture Facility |  |
| Cell line | Raji B cells | UCSF Cell Culture Facility |  |
| Plasmid | CAR (1<sup>st</sup>)-sfGFP | This study | Signal peptide: CD8a (Gene Bank AAH25715 aa 1-21), extracellular antibody: anti-CD19 CAR (Gene Bank AMZ04819, aa 23-130-linker (3xGGGGS)- aa 148-267), Stalk and Transmembrane: CD8a (Gene Bank AAH25715 aa 138-206), Cytosolic: CD3z (Gene Bank AAH25703 aa 52-164), linker (2xGGS), sfGFP (Gene Bank AVR55105) |
| Plasmid | CAR (3<sup>rd</sup>)-sfGFP | This study | Signal peptide: CD8a (Gene Bank AAH25715 aa 1-21), extracellular antibody: anti-CD19 CAR (Gene Bank AMZ04819, aa 23-130-linker (3xGGGGS)- aa 148-267), Stalk and Transmembrane: CD8a (Gene Bank AAH25715 aa 138-206), Cytosolic: CD28 (Gene Bank AAA51944 aa 180-220) -4-1BB (Gene Bank AAA53133 aa 214-255)-CD3z (Gene Bank AAH25703 aa 52-164), linker (2xGGS), sfGFP (Gene Bank AVR55105) |
| Plasmid | Gads-mCherry | This study | FKBP (Gene Bank AAP36774 aa 1-109), linker (3xGS-G4S2), Gads (AAH26002 aa 1-330), linker (2xGGS), mCherry (AMO27233 aa 1-236) |
| Protein | CD19 (Ex), biotinylated | Sino biological | A DNA sequence encoding CD19 (UniProt P15391) aa 1-291 was expressed with a C-terminal polyhistidine tag in HEK293 cells. The purified protein was biotinylated in vitro. |
| Protein | ICAM-1(Ex)-His | Sino biological | A DNA sequence encoding ICAM1 (NP_000192.2) aa 1-480 was fused with the a polyhistidine tag at the C-terminus. The gene product was expressed and purified from HEK293 cells. |
| Antibody | Anti-LAT antibody | Cell Signaling | Host: Rabbit. Cat #9166s. |
| Antibody | Anti-p150 (glued) antibody | BD Biosciences | Host: Mouse. Cat #612709. |
| Antibody | IRDye 800<br>CW goat anti-rabbit | Li-Cor<br>Biosciences | Cat #926-32211. |
| Antibody | IRDye<br>800CWGoat<br>anti-mouse | Li-Cor<br>Biosciences | Cat #924-32210. |
| Antibody | Anti-pSLP76<br>antibody | Abcam | Rabbit monoclonal [EP2853Y] to SLP76<br>(phospho Y145). Cat #ab75829. |
| Antibody | Goat anti-Rabbit A647 | Molecular<br>Probes | Goat anti-Rabbit IgG (H+L) Cross-Adsorbed<br>Secondary Antibody, Alexa Fluor 647. Cat # A-21244 |
| Antibody | Biotin-OKT3 | BioLegend | Biotin anti-human CD3 Antibody, Cat # 317319 |

### Cell Culture

Jurkat T cell lines were grown in RPMI (Invitrogen) supplemented with 10% FBS (Invitrogen), 1% PenStrep-Glutamine, and 10 mM HEPES pH 7.4. HEK293T cells (purchased from the ATCC collection) were grown in DMEM (Invitrogen) supplemented with 10% FBS and 1% PenStrep-Glutamine.

### Lentiviral Product and Generation of Stable Expressing Jurkat Cell Lines

HEK293T cells were co-transfected with the pHR transfer plasmids with second generation packaging plasmids pMD2.G and psPAX2 (Addgene plasmid #12259 and #12260) using Lipofectamine LTX Reagent (ThermoFisher). 48 hr after plasmid transfection, cell culture media containing virus particles were harvested, centrifuged and filtered through 0.45 μm pore size filter, and mixed with Jurkat cells for infection in RPMI media for 72 hr. Jurkat cells expressing fluorescent proteins were FACS sorted to generate a stable and homogenous expression population.

### Primary T Cell Gene Editing

T cell editing was performed according to published protocols^21,22^. Briefly, peripheral blood mononuclear cells (PBMCs) were isolated from healthy donors (Trima Apheresis, Blood Centers of the Pacific) by Ficoll centrifugation with SepMate tubes (STEMCELL Technologies). Bulk T cells were isolated from PBMCs using Easysep Human T Cell Isolation Kit (STEMCELL Technologies) and activated with Dynabeads Human T-Activator CD3/CD28 (Thermo Fisher Scientific) at a 1:1 (bead:cell) ratio. Cells were cultured in X-VIVO 15 medium (Lonza Bioscience) supplemented with 10% fetal bovine serum (FBS), 50 μM 2-mercaptoethanol (Thermo Fisher Scientific), 10 mM N-acetyl L-cystine (Sigma), 500 U/ml Interleukin-2 (IL-2, Aldesleukin Proleukin, Novartis), 5 ng/ml IL-7 and 5 ng/ml IL-15 (R&D Systems).

24 hr after activation, 10 μL of concentrated lentivirus was added per ml of T cells. Lentivirus was produced as previously described^22^. Briefly, 293T cells were transfected with transfer plasmid, pMD2.G and psPAX2 (Addgene) using FugeneHD transfection reagent (Promega Corporation). 24 hr later, Viral Boost reagent #VB100 (Alstem) was added according to the manufacturer’s protocol. Viral supernatant was collected 48 hr after transfection and concentrated 100x using lentivirus precipitation solution #VC100 (Alstem).

48 hr after transduction, culture was washed, Dynabeads were removed and T cells were counted. Ribonucleoproteins (RNPs) were produced as previously described (Roth et al., Nature 2018). Briefly, crRNA and tracrRNA (Dharmacon) were resuspended at a concentration of 160 μM in nuclease-free duplex buffer (IDT), mixed 1:1 by volume and incubated for 30 min at 37° C to form the gRNA. Poly-L-glutamic acid (PGA) sodium salt (Sigma, stock 100 mg/ml) and recombinant Cas9 (Macrolab UC, stock 40 μM) were added as previously described (0.72:1 by volume for PGA vs gRNA and 2:1 by volume for Cas9 vs gRNA)^23^ and incubated at 37° C for 15 min to form RNP complex.

Prior to electroporation, T cells were centrifuged for 10 min at 90x g and resuspended in electroporation buffer P3 (Lonza) at a concentration of 750,000 cells/ 20 μL. 750,000 cells per well were mixed with 6 μL of RNP per well and electroporated in a 96-well electroporation plate (Lonza) using electroporation code EH115 on a 4D Nucleofector system with 96-well shuttle (Lonza). Immediately after electroporation, 80 μL of pre-warmed media without cytokines were added per well and cells were incubated in the electroporation plate at 37° C. After 15 min, cells were moved to final culture vessels. Following electroporation, T cells were cultured in media without IL-7 and −15 (only IL-2) and split every two days.

LAT gRNA sequence: TTTACCAGTTTGTATCCAAG (from Brunello sgRNA library, Doench et al., Nature Biotechnology 2016).

psPAX2 was a gift from Didier Trono (Addgene plasmid # 12260; http://n2t.net/addgene:12260; RRID:Addgene_12260). pMD2.G was a gift from Didier Trono (Addgene plasmid # 12259; http://n2t.net/addgene:12259; RRID:Addgene_12259).

### Preparation of Supported Lipid Bilayer

Lipid mixture were prepared with 97.5% POPC, 2.0% DGS-NGA-Ni, 0.5% PEG5000-PE, and less than 0.1% Biotin-Cap-PE. Lipids were dissolved in chloroform in glass tubes, and dried under a stream of argon gas followed by further drying in the vacuum for 2 hr. The dried lipid films were then hydrated with PBS pH 7.4 (invitrogen). The small unilamellar vesicles (SUVs) were produced by twenty freeze-thaw cycles (−80°C and 37°C) and collected as the supernatant after centrifuge at 53,000x g for 45 min at 4°C. SUVs were stored at 4°C and used within 2 weeks.

Glass coverslip (Ibidi Cat#10812) were RCA-cleaned followed by extensive washing with pure water, and dried with nitrogen. PDMS (Dow Corning) wells were made by preparing PDMS substrate mixtures according to the manufacturer’s instructions and casting the PDMS mixtures into laser-cut acrylic mold. To build supported lipid bilayer, PDMS wells and glass coverslips were cleaned with plasma in a Harrick Plasma cleaner before assembling them into glass-bottomed PDMS chambers. SUV suspensions were then deposited in each chamber and allows to form for 1 hr. After 1 hr, wells were washed extensively with PBS, and functionalized by incubation for 1 hr with proteins as designated.

All the following lipids were purchased from Avanti Polar Lipids: 16:0-18:1 POPC 1-palmitoyl-2-oleoyl-sn-glycero-3-phosphocholine (POPC; Cat #850457), 18:0 PEG5000 PE 1,2-distearoyl-sn-glycero-3-phosphoethanolamine-N-[methoxy(polyethylene glycol)-5000] (ammonium salt) (PE-PEG5000; Cat #880220), 18:1 DGS-NTA (Ni+2) 1,2-dioleoyl-sn-glycero-3-[(N-(5-amino-1-carboxypentyl)iminodiacetic acid)succinyl] (nickel salt) (Ni2+-NTA-DOGS; Cat#790404), 1,2-dipalmitoyl-sn-glycero-3-phosphoethanolamine-N-(cap biotinyl) (sodium salt) (Biotin-Cap-PE; Cat #870277).

### IL-2 Production

Raji B cells were co-cultured at a 1:1 ratio with Jurkat CAR T cells in RPMI medium supplemented with 20 mM HEPES-Na^+^, pH7.4 for 24 hours at 37°C. Supernatant was collected for IL-2 measurement using an ELISA kit (BioLegend #431801).

### Cell Conjugation Assay

Raji B or K562 cells expressing a membrane marker mCherry-CAAX were co-cultured at a 1:1 ratio with Jurkat CAR T cells in RPMI medium supplemented with 20 mM HEPES-Na^+^, pH7.4 for 1 hr at 37°C. Cells were fixed in 1.6% paraformaldehyde for 15 min at RT, washed, and resuspended in RPMI medium supplemented with 20 mM HEPES and imaged by confocal microscopy.

### Imaging Setup

Imaging was performed on a Nikon TI-E microscope equipped with a Nikon 100× Plan Apo 1.49 NA oil immersion objective, four laser lines (405, 488, 561, and 640 nm), a Hamamatsu Flash 4.0, and μManager software. A polarizing filter was placed in the excitation laser path to polarize the light perpendicular to the plane of incidence. The angle of illumination was controlled with either a standard Nikon TIRF motorized positioner or a mirror moved by a motorized actuator (CMA-25CCCL; Newport). Data collection was performed at 37°C for the experiments involving live cells, and at room temperature for the rest non-cell experiments. Before imaging, cells were pelleted, washed, and resuspended in the imaging buffer containing 20 mM HEPES pH 7.4, 1 mM CaCl_2_, 135 mM NaCl, 0.5 mM MgCl_2_, 4 mM KCl, and 10 mM glucose.

### Image Analysis

Images were analyzed using Fiji. The same brightness and contrast were applied to images within the same panels. To quantify the clustering levels (as shown in Figure 2B and 3B), a uniform cell-sized circular region of interest (ROI) that is of 10 μm in diameter was manually placed over the region of cell fluorescence. The average and the standard deviation of fluorescence intensity inside the ROI was measured respectively, and the normalized variance, i.e. the ratio of the standard deviation fluorescence intensity divided by the average fluorescence intensity, was used to indicate the dispersive distribution of fluorescence intensity for each cell. To quantify pSLP76 levels by immunofluorescence (Figure 3B and 4C) or Gads recruitment (Figure 3A), a uniform cell-sized circular ROI that is of 10 μm in diameter was manually placed over the region of cell fluorescence, and the fluorescence intensity of the channel of interest inside that ROI was measured.

### Immunoblotting

Cells were washed twice in cold PBS and lysed on ice for 15 min in lysis buffer containing 50 mM Tris, 150 mM NaCl, 0.1% (w/v) SDS, 0.5% (w/v) sodium deoxycholate, 1% (v/v) TritonX-100, pH 7.5 supplemented with protease inhibitor cocktail (Roche) and benzonase (Novagen). Cell lysates were then centrifuged at 21,000 xg for 20 min at 4°C. The supernatants were processed for SDS-PAGE and immunoblotting with standard procedure. Immunoblots were visualized using Odyssey Fc Imager (LI-COR Biosciences) at 800 nm channel, and quantified using the gel analysis function in Fiji.

### Immunofluorescence

Cells were activated on supported lipid bilayers in the imaging buffer (see section “Imaging Setup” for formula) for 30 min at 37°C, and fixed by adding PFA to a final concentration of 3.2% (v/v) and incubating at room temperature for 15 min. Cells were then washed twice with PBS, permeabilized with methanol at 4°C for 10 min. Then cells were blocked with PBS containing 0.1% (v/v) tween20, 1% (w/v) BSA, and 22.5 mg/mL glycine for one hour, and incubated with the primary antibody in the same buffer overnight. The next day, cells were washed three times with PBS, incubated with buffer containing Alexa-647 conjugated secondary antibody for 1 hour. Excess antibodies were washed away before proceeding to image acquisition.

## References

1 Ghobadi, A. Chimeric antigen receptor T cell therapy for non-Hodgkin lymphoma. Curr Res Transl Med 66, 43–49, doi:10.1016/j.retram.2018.03.005 (2018).

2 Lee, D. W. et al. T cells expressing CD19 chimeric antigen receptors for acute lymphoblastic leukaemia in children and young adults: a phase 1 dose-escalation trial. Lancet 385, 517–528, doi:10.1016/S0140-6736(14)61403-3 (2015).

3 Porter, D. L., Levine, B. L., Kalos, M., Bagg, A. & June, C. H. Chimeric antigen receptor-modified T cells in chronic lymphoid leukemia. N Engl J Med 365, 725–733, doi:10.1056/NEJMoa1103849 (2011).

4 June, C. H. & Sadelain, M. Chimeric Antigen Receptor Therapy. N Engl J Med 379, 64–73, doi:10.1056/NEJMra1706169 (2018).

5 Lim, W. A. & June, C. H. The Principles of Engineering Immune Cells to Treat Cancer. Cell 168, 724–740, doi:10.1016/j.cell.2017.01.016 (2017).

6 Davenport, A. J. et al. Chimeric antigen receptor T cells form nonclassical and potent immune synapses driving rapid cytotoxicity. Proceedings of the National Academy of Sciences of the United States of America 115, E2068–E2076, doi:10.1073/pnas.1716266115 (2018).

7 Xhangolli, I. et al. Single-cell Analysis of CAR-T Cell Activation Reveals A Mixed TH1/TH2 Response Independent of Differentiation. Genomics Proteomics Bioinformatics 17, 129–139, doi:10.1016/j.gpb.2019.03.002 (2019).

8 Valvo, S. et al. Comprehensive Analysis of Immunological Synapse Phenotypes Using Supported Lipid Bilayers. Methods in molecular biology 1584, 423–441, doi:10.1007/978-1-4939-6881-7_26 (2017).

9 Bunnell, S. C. et al. T cell receptor ligation induces the formation of dynamically regulated signaling assemblies. J Cell Biol 158, 1263–1275, doi:10.1083/jcb.200203043 (2002).

10 Varma, R., Campi, G., Yokosuka, T., Saito, T. & Dustin, M. L. T cell receptor-proximal signals are sustained in peripheral microclusters and terminated in the central supramolecular activation cluster. Immunity 25, 117–127, doi:10.1016/j.immuni.2006.04.010 (2006).

11 Grakoui, A. et al. The immunological synapse: a molecular machine controlling T cell activation. Science 285, 221–227 (1999).

12 Zhang, W., Sloan-Lancaster, J., Kitchen, J., Trible, R. P. & Samelson, L. E. LAT: the ZAP-70 tyrosine kinase substrate that links T cell receptor to cellular activation. Cell 92, 83–92, doi:10.1016/s0092-8674(00)80901-0 (1998).

13 Su, X. et al. Phase separation of signaling molecules promotes T cell receptor signal transduction. Science 352, 595–599, doi:10.1126/science.aad9964 (2016).

14 Barda-Saad, M. et al. Dynamic molecular interactions linking the T cell antigen receptor to the actin cytoskeleton. Nat Immunol 6, 80–89, doi:10.1038/ni1143 (2005).

15 Finco, T. S., Kadlecek, T., Zhang, W., Samelson, L. E. & Weiss, A. LAT is required for TCR-mediated activation of PLCgamma1 and the Ras pathway. Immunity 9, 617–626, doi:10.1016/s1074-7613(00)80659-7 (1998).

16 Bunnell, S. C., Kapoor, V., Trible, R. P., Zhang, W. & Samelson, L. E. Dynamic actin polymerization drives T cell receptor-induced spreading: a role for the signal transduction adaptor LAT. Immunity 14, 315–329, doi:10.1016/s1074-7613(01)00112-1 (2001).

17 Liu, S. K., Fang, N., Koretzky, G. A. & McGlade, C. J. The hematopoietic-specific adaptor protein gads functions in T-cell signaling via interactions with the SLP-76 and LAT adaptors. Curr Biol 9, 67–75, doi:10.1016/s0960-9822(99)80017-7 (1999).

18 Wunderlich, L., Farago, A., Downward, J. & Buday, L. Association of Nck with tyrosine-phosphorylated SLP-76 in activated T lymphocytes. Eur J Immunol 29, 1068–1075, doi:10.1002/(SICI)1521-4141(199904)29:04<1068::AID-IMMU1068>3.0.CO;2-P (1999).

19 Oancea, E. & Meyer, T. Protein kinase C as a molecular machine for decoding calcium and diacylglycerol signals. Cell 95, 307–318 (1998).

20 Bartelt, R. R., Cruz-Orcutt, N., Collins, M. & Houtman, J. C. Comparison of T cell receptor-induced proximal signaling and downstream functions in immortalized and primary T cells. PLoS One 4, e5430, doi:10.1371/journal.pone.0005430 (2009).

21 Schumann, K. et al. Generation of knock-in primary human T cells using Cas9 ribonucleoproteins. Proc Natl Acad Sci U S A 112, 10437–10442, doi:10.1073/pnas.1512503112 (2015).

22 Shifrut, E. et al. Genome-wide CRISPR Screens in Primary Human T Cells Reveal Key Regulators of Immune Function. Cell 175, 1958–1971 e1915, doi:10.1016/j.cell.2018.10.024 (2018).

23 Nguyen, D. N. et al. Polymer-stabilized Cas9 nanoparticles and modified repair templates increase genome editing efficiency. Nat Biotechnol, doi:10.1038/s41587-019-0325-6 (2019).

